# Single-Cell Transcriptomic Mapping of PD-L1/TLR4 Remodeling Informs Topical Immunoprevention Timing in Skin Carcinogenesis

**DOI:** 10.64898/2026.06.01.729458

**Authors:** Ran Wei, Rudramani Pokhrel, Delaney Stratton, Sara Centuori, Clara Curiel-Lewandrowski, Georg T. Wondrak, Sally E. Dickinson, Bonnie J. LaFleur, Xiaoxiao Sun

**Author notes:** These authors contributed equally to this work and share first authorship. Corresponding authors: Correspondence should be addressed to: Bonnie J. LaFleur, PhD, R. Ken Coit College of Pharmacy The University of Arizona Tucson, Arizona, USA, Email: [ ] Phone: [], Xiaoxiao Sun, PhD, Department of Epidemiology and Biostatistics, Mel and Enid Zuckerman College of Public Health The University of Arizona, Tucson, Arizona, USA, Email: [ ] Phone: [].

## Abstract

Cutaneous squamous cell carcinoma (cSCC) represents a growing public health burden, with incidence projected to increase 23–29% over the coming decade. Topical immunoprevention strategies targeting the `PD-L1`/`PD-1` and `TLR4` axes have demonstrated preclinical efficacy, yet optimal intervention timing in humans remains undefined. To address this gap, single-cell RNA sequencing was performed on matched sun-protected (SP), sun-damaged (SD), and actinic keratosis (AK) biopsies from the same individuals, along with independent cSCC cases. Immune checkpoint and innate inflammatory signals were detectable as early as SD skin, prior to histologically confirmed dysplasia. Monotonically increasing expression of `CD274` (`PD-L1`), `CTLA4`, `PDCD1`, `CD27`, and `STAT1`, alongside progressive `TLR4`–`MYD88` innate immune signaling, was revealed through pseudobulk data analysis, with earliest upregulation at the SD stage. Fuzzy c-means trajectory clustering identified cell-typespecific programs across dendritic cells, macrophages, T cells, fibroblasts, endothelial cells, and keratinocytes. Dendritic cells shifted from early inflammatory antigen-presenting programs toward late `PD-L1`/`IFN`-regulatory states; macrophages showed monotonically increasing `TLR4`-associated myeloid activation; and T cells defined a “hot but exhausted” microenvironment in established cSCC. These findings identify SD and AK as biologically active stages for topical immunoprevention and provide a cellular roadmap for `PD-L1`/`PD-1` and `TLR4` blockade strategies.

## 1 Introduction

Cutaneous squamous cell carcinoma (cSCC) is one of the most common malignancies worldwide and represents a major and growing public health burden [Tokez et al., 2020]. In the United States, cSCC represents 20-25% of all keratinocytic neoplasms, which could result in ∼900,000 new SCC cases annually, with recent European statistics estimating a 23-29% increase in rates between 2017 and 2027 [Rogers et al., 2015]. The average annual total cost of skin cancer treatment in the United States increased from $3.6 to $8.1 billion between 2002 and 2011 [Guy Jr et al., 2015], and surgical procedures account for the majority of expenditures [Chen et al., 2016]. This burden is disproportionately high in sun-exposed populations, including residents of the American Southwest, where rates approximate those reported in Australia. With increasing life expectancy and the associated decline in immune surveillance known as immunosenescence, the societal and healthcare burden of nonmelanoma skin cancers (NMSC) is projected to worsen substantially unless effective targeted prevention strategies are implemented.

The development of cSCC follows a well-characterized multistep carcinogenesis sequence initiated by chronic exposure to ultraviolet (UV) radiation. UV-induced DNA damage and immune dysregulation in keratinocytes drive the transition from normal sun-protected skin (SP) to sun-damaged skin (SD) and actinic keratosis (AK), a precancerous lesion of epidermal keratinocytic dysplasia, to invasive cSCC. AKs represent a clinically accessible and biologically important intervention point. Although individual AKs carry a relatively low short-term malignant transformation risk, patients with skin malignancy and high AK burden face a substantially elevated cumulative risk of progression to invasive disease [Criscione et al., 2009]. An effective intervention at this stage or earlier could significantly reduce the incidence of invasive cSCC and the associated procedural burden. The carcinogenic potential of chronically sun-exposed skin is further amplified by the biology of aging, as cSCC is often associated with years of UV-exposure prior to malignancy. Furman et al. recently synthesized evidence that UV-induced genomic instability, inflammation, and accumulation of senescent cells in the skin are tightly interconnected hallmarks that collectively accelerate the transition from normal tissue homeostasis to a pro-carcinogenic microenvironment [Furman et al., 2025]. These processes associated with aging are active contributors to the immune dysregulation that characterizes the SP-cSCC trajectory investigated here.

Preclinical evidence from UV-induced murine skin tumor models has established two immune pathways as particularly promising targets for topical immunoprevention, specifically therapeutic agents targeting the `PD-L1`/`PD-1` immune checkpoint axis and `TLR4`-associated innate inflammatory signaling. Topical small-molecule `PD-L1` blockade has been shown to inhibit UV-induced stress signaling and inflammatory responses in SKH-1 mouse skin [Dickinson et al., 2024], while pharmacological `TLR4` antagonism using resatorvid has been shown to block solar UV-induced skin tumorigenesis in the same model [Blohm-Mangone et al., 2018]. These findings have motivated ongoing efforts to translate these strategies into human topical immunoprevention approaches, such as the prior work identifying `PD-1`/`PD-L1` axis as an emerging therapeutic and prevention target in keratinocytic skin cancer [Vaishampayan et al., 2023]. However, a critical translational gap remains: the stage at which `PD-L1`/`PD-1` and `TLR4` pathways become biologically active during human skin carcinogenesis is not known. Defining this timing is essential for identifying the optimal window for topical immunoprevention.

Single-cell RNA sequencing (scRNA-seq) has transformed the study of the tumor microenvironment by enabling simultaneous transcriptomic profiling of cell types present in a tissue at single-cell resolution.The use of spatial transcriptomics and scRNA-seq in dermatological research has grown substantially in recent years, including healthy skin characterization, wound healing, inflammatory conditions, and cutaneous malignancies, including cSCC [Schepps et al., 2024]. Previous spatial transcriptomic work showed that dermal transcriptional changes occur during progression from sun-protected skin to actinic keratosis, identifying the dermal compartment as a key site of early transcriptional remodeling [LaFleur et al., 2023]. Recent scRNA-seq studies in cSCC and related keratinocytic malignancies have revealed substantial heterogeneity in both epithelial and immune compartments and have characterized the immune landscape of AK and early cSCC [Xu et al., 2025, Zou et al., 2023]. However, these studies have generally not been designed to evaluate immunoprevention strategies specifically. The present study enables the resolution of specific immune cell populations and pathway dynamics associated with `PD-L1`/`PD-1` and `TLR4` pathways.

Importantly, it also defines the stages at which these cellular and molecular changes emerge during skin carcinogenesis. The experiment uses an annotated cohort of matched SP, SD, and AK biopsies together with independent cSCC cases. We apply pseudobulk differential expression analysis, pathway enrichment, and targeted fuzzy c-means trajectory clustering to map the stage-specific dynamics of immune checkpoint and innate inflammatory pathways in cell types present in human skin. Our findings identify SD and AK skin as biologically active and clinically accessible stages for topical immunoprevention and provide a cellular and molecular roadmap to guide the selection and timing of `PD-L1`/`PD-1` and `TLR4` blockade strategies for the prevention of cSCC.

## 2 Results

### 2.1 Cellular composition shifts from a structural skin landscape toward an immune-enriched tumor microenvironment

Uniform Manifold Approximation and Projection (UMAP) visualization showed the expected diversity of epithelial, stromal, vascular, and immune populations, including keratinocytes, fibroblasts, endothelial cells, macrophages, dendritic cells, T cells, NK cells, mast cells, melanocytes, plasma/B cells, and several smaller accessory populations (Figure 1a). Cell-type proportions varied across tissue states (Figure 1b). cSCC samples were enriched for immune populations, particularly T cells, macrophages, NK cells, mast cells, plasma/B cells, and neutrophils. By contrast, SP and SD samples were dominated by structural compartments, especially endothelial cells and fibroblasts, whereas AK showed an intermediate composition with persistent stromal representation and increasing immune-cell abundance (Figure 1c). Dendritic cells were slightly more abundant in cSCC, while endothelial cells, fibroblasts, and keratinocytes shifted only slightly across states. Taken together, these descriptive cell-level summaries describe a transition from a predominantly structural skin microenvironment in SP and SD tissue toward an immune-enriched and tumorassociated microenvironment in cSCC, consistent with prior single-cell studies and recent reviews of AK-to-cSCC progression and non-melanoma skin cancer (NMSC) [Zou et al., 2023, Li et al., 2025, Mousa et al., 2024, Yan et al., 2024].

**Figure 1.**
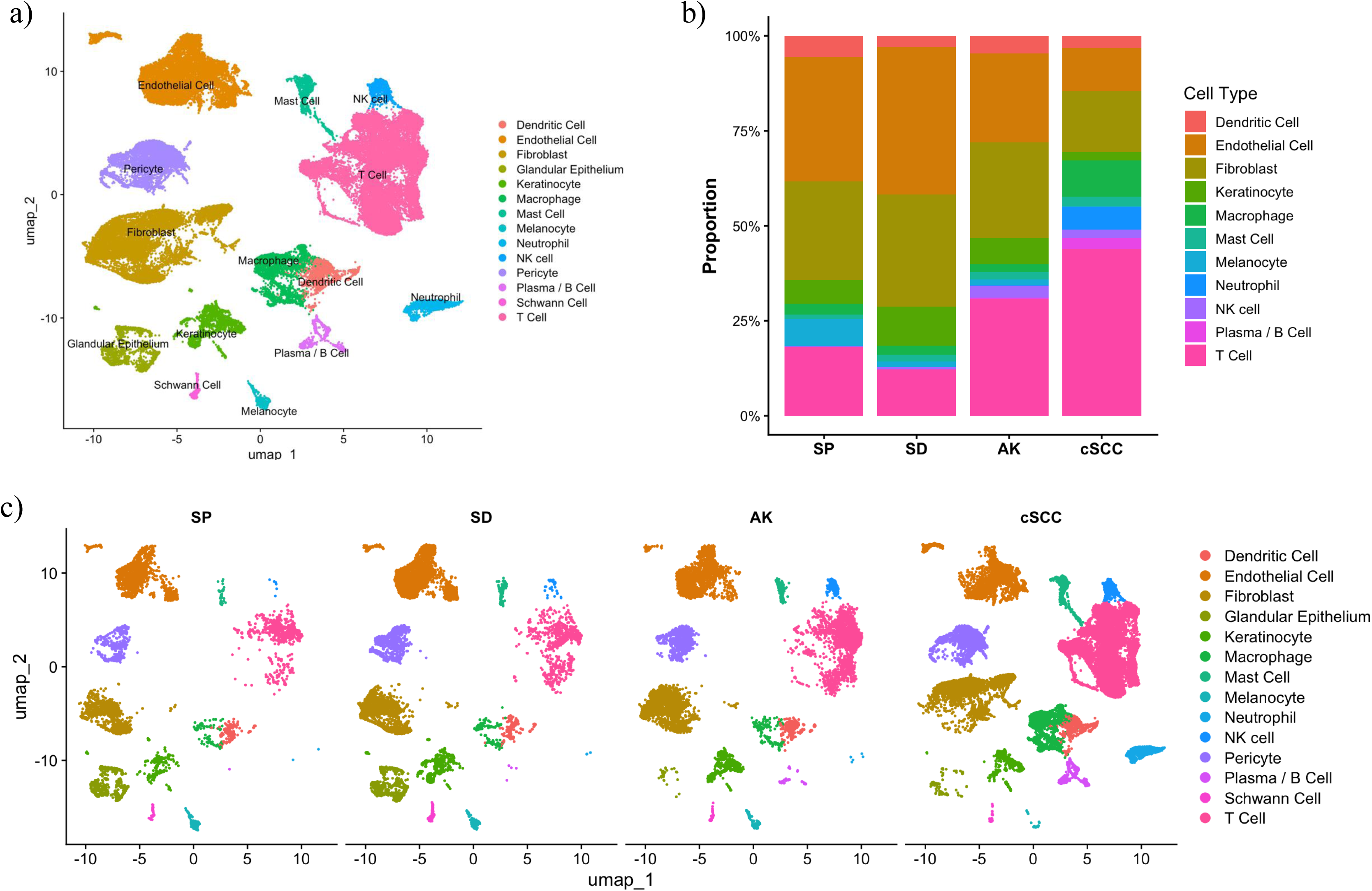
Single-cell transcriptomic landscape across disease progression from sun-protected to cutaneous squamous cell carcinoma. a) UMAP visualization of integrated single-cell RNA sequencing data showing major cell populations identified across all samples. b) Stacked bar plots showing the relative proportions of major cell populations across sun-protected skin (SP), sun-damaged skin (SD), actinic keratosis (AK), and cutaneous squamous cell carcinoma (cSCC). Progressive enrichment of immune and stromal cell populations and small shifts in keratinocyte composition were observed during disease progression. c) UMAP plots stratified by disease progression demonstrating dynamic changes in cellular composition and transcriptional states from SP and SD through AK to cSCC.

### 2.2 Progressive dysregulation of PD-L1 and TLR4 pathways during disease development

Having established the overall compositional landscape, we next examined the expression of genes specifically from the `PD-L1`/`PD-1` and `TLR4` signaling pathways (Figure 2). This analysis was performed across all conditions using pseudobulk log-CPM normalized expression, enabling stagewise comparisons that account for between-individual variability. Among `PD-L1` pathway-associated genes, `CD274` (`PD-L1`), `PDCD1` (`PD-1`), `CTLA4`, `CD27`, and `CD70` all demonstrated statistically significant increasing expression across the SP–SD–AK–cSCC continuum, with BenjaminiHochberg corrected p-values (*p*_adj_) ranging from 0.003 to 0.015. The earliest detectable upregulation for several of these genes was observed at the SD stage, supporting the hypothesis that immune checkpoint engagement is initiated prior to histologically confirmed dysplasia. `STAT1`, a transcription factor downstream of both `IFN` signaling and `TLR4`-associated pathways, was also significantly upregulated (*p*_adj_ = 0.003), indicating convergent activation of both innate and adaptive immune signaling axes during carcinogenesis. Upregulation of `STAT1` and `CD274` suggests a coordinated interferon-driven transcriptional program, consistent with `STAT1`-mediated induction of `PD-L1` as lesions progress [Garcia-Diaz et al., 2017] into an immune landscape that accompanies cSCC progression [Bailey et al., 2023]. `STAT1` and `CTLA4` are monotonically increasing (Jonckheere–Terpstra *p*_adj_ = 0.003). This aligns with recent evidence in cSCC, as possible evidence of `STAT` acting as a dual regulator, both as a biomarker of active `IFN`-*γ*-driven anti-tumor surveillance and a key mechanistic driver of immune evasion via checkpoint induction [Jiang et al., 2024].

**Figure 2.**
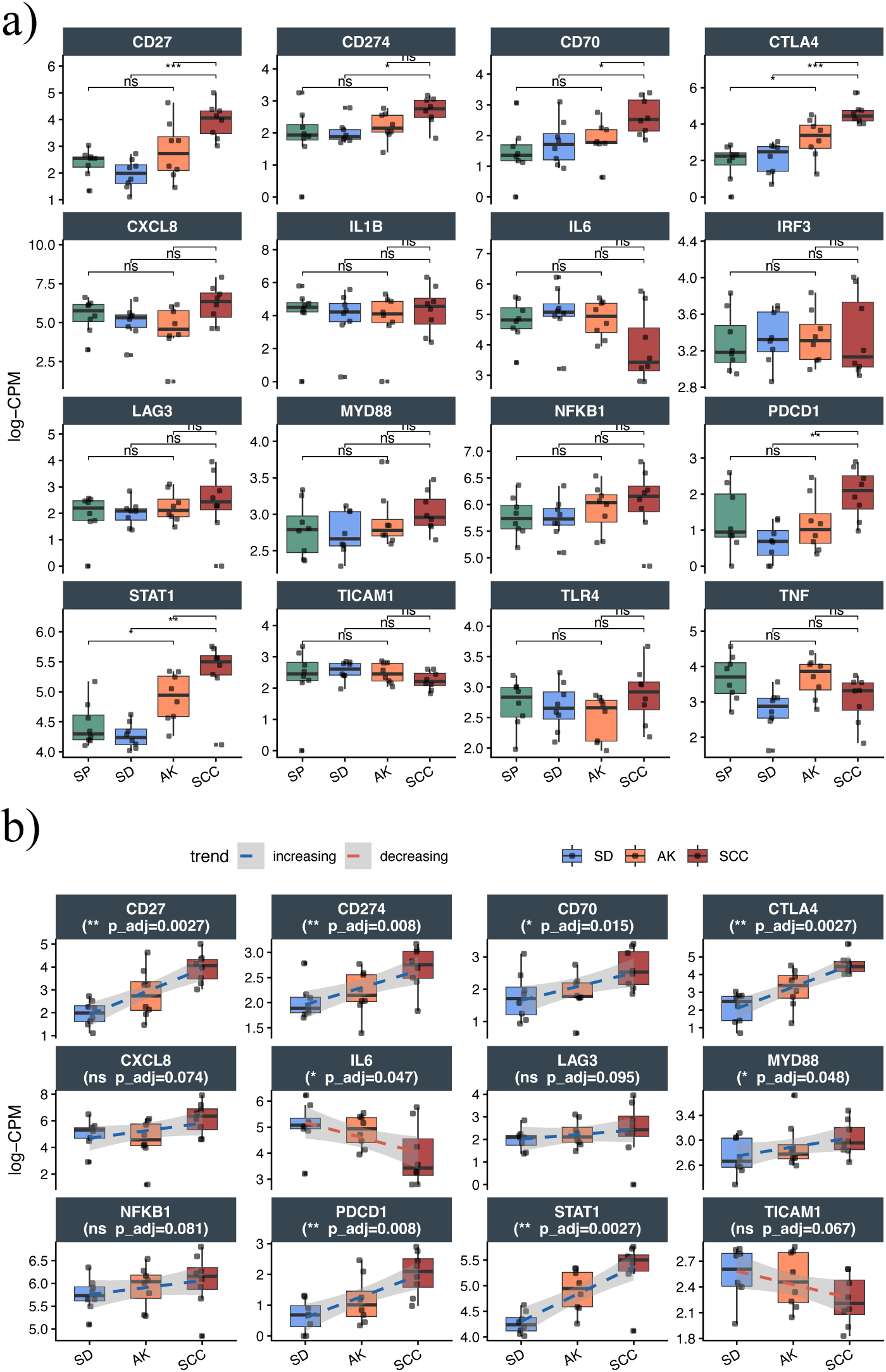
PD-L1 and TLR4 signaling–related gene expression from sun-protected skin to cutaneous squamous cell carcinoma. a) Box plots showing normalized expression levels (log-CPM) of `PD-L1` and `TLR4` signaling–related genes across SP, SD, AK, and cSCC. b) Trend analysis of selected genes demonstrating progressive increases or decreases in expression from SD through AK to cSCC. Abbreviations: SP, sun-protected skin; SD, sun-damaged skin; AK, actinic keratosis; cSCC, cutaneous squamous cell carcinoma.

Within the `TLR4` signaling pathway, `MYD88` showed a statistically significant monotonically increasing trend (*p*_adj_ = 0.048), consistent with activation of canonical `TLR4`–`MyD88`-dependent innate inflammatory signaling typically shown in development of UV-driven diseases [Salcedo et al., 2013, Sherwani et al., 2025]. Additionally, there was upregulation of `NFKB1` (*p*_adj_ = 0.081) and `CXCL8` (*p*_adj_ = 0.074), both previously shown to be canonical downstream effectors of `MYD88`-dependent NF-*κ*B activation. These results further support progressively amplified innate inflammatory responses across the malignancy trajectory[De Falco et al., 2024, Tuong et al., 2019]. In particular, NF-*κ*B activation downstream of `MYD88` has been shown to contribute to `CD274` (`PD-L1`) transcriptional induction, suggesting that `TLR4`–`MyD88` signaling may synergize with the `IFN`-*γ*-`STAT1` axis to reinforce the immune-evasive landscape in the progression of cSCC lesions [Antonangeli et al., 2020, Yin et al., 2021, Dickinson and Wondrak, 2019].

Not all pathway genes showed monotonic increases. `IL6` exhibited a monotonically decreasing trend across tissue subtypes (*p*_adj_ = 0.047 and 0.067, respectively), and while `TICAM1`showed a similar decreasing pattern, it was not statistically significant after p-value correction (*p*_adj_ = 0.067). Although `IL6` has been implicated in promoting early skin SCC growth [Lederle et al., 2011], its decline in established cSCC may reflect a transition from acute inflammatory signaling toward a more chronic, tolerogenic microenvironment that is consistent with progressive immunoediting [Ciąyżńska et al., 2021]. Similarly, the attenuation of `TICAM1` suggests selective rewiring of innate immune signaling away from the immunostimulatory type I interferon branch. Together, these divergent trends indicate that while upstream innate sensing is progressively amplified along the `TLR4`–`MyD88` axis, certain downstream effector cell populations are simultaneously redirected; this is a pattern consistent with the emergence of an immunosuppressive microenvironment in established cSCC. Collectively, these findings demonstrate that progressive dysregulation of both the `PD-L1` / `PD-1` checkpoint axis and innate immune signaling of `TLR4`–`MyD88` is a defining feature of NMSC, detectable as early as the SD stage and amplified through AK into established cSCC. Together, these data support a model in which the tumor microenvironment of cSCC transitions from an innate-dominant inflammatory state in SD skin to a checkpoint-dominated immunosuppressive landscape consistent with the phenotype “hot but exhausted” that underlies the demonstrated clinical responsiveness of advanced cSCC to `PD-1` blockade observed with cemiplimab [Migden et al., 2018].

### 2.3 Trajectory analysis of gene clusters within PD-L1/PD-1 and TLR4 pathways

The targeted fuzzy c-means clustering analysis revealed coordinated stage-specific remodeling across six cells compartments: dendritic cells, macrophages, T cells, fibroblasts, endothelial cells, and keratinocytes (Fugre 3). The targeted gene panel represented six functional categories: `PD-L1` / `PD-1` checkpoint regulation, `TLR4`-associated innate inflammatory signaling, IFN/JAK–STAT- mediated `PD-L1` induction, antigen presentation/APC maturation, and immunoregulatory/tissue-remodeling support. Trajectory analyses were performed for cell types with sufficient representation in all four stages of disease (subtypes). Macrophages were sparsely represented in SP, SD, and AK but expanded substantially in cSCC (Figures 1b and 1c); consequently, only one group of macrophages yielded interpretable gene-level trajectories. Optimal fuzzification parameters, *f*, were used for all genes within a cell population. These were, 3.45 (dendritic cells); 3.32 (endothelial cells); 3.27 (fibroblasts); 3.82 (macrophages); 3.34 (T cells); and 4.82 (keratinocytes).

#### TLR4-associated myeloid activation

The clearest `TLR4` signal emerged from macrophages. A single interpretable macrophage cluster (cluster 2) increased progressively from SP through cSCC and contained `CD14` and `LY96`, the two core components of the `TLR4` receptor complex, together with the costimulatory marker `CD86` and the tolerogenic chemokine `CCL18`. This co-expression pattern is consistent with a mixed M1/M2-like tumor-associated macrophage state and with progressive TLR4 activation accumulating throughout carcinogenesis rather than arising de novo in established tumors. A complementary signal was observed in fibroblasts: cluster 1 increased progressively from SP to cSCC and contained `HMGB1`, `IL6`, `STAT1`, `CXCL9`, and `CXCL10`, consistent with an accumulating IFN-responsive stromal inflammatory program. The co-upregulation of `HMGB1` in fibroblasts alongside `LY96` in macrophages is consistent with a self-sustaining DAMP-driven cycle linking UV-induced tissue damage to chronic innate inflammatory signaling. A second fibroblast cluster showed a sharp late rise into cSCC with `LY96`, `IDO1`, `MX1`, `TAP1`, and `CXCL13`, suggesting that stromal immunoregulatory and antigen-presentation-associated features intensify at the invasive stage. A third fibroblast cluster declined toward cSCC and contained TLR adaptor genes `TICAM2`, `IRAK4`, and `TRAF6`, suggesting upstream innate signaling components are redistributed or replaced as disease progresses.

#### PD-L1/IFN remodeling in dendritic cells

Three dendritic cell trajectory clusters captured a shift from early inflammatory antigen-presenting programs toward late immunoregulatory states. Cluster 3 was elevated in SP and SD before declining into cSCC and contained pro-inflammatory mediators `IL1B`, `TNF`, `CLEC10A`, and `CXCL8`, consistent with an early resident APC program that is reduced or functionally replaced during malignant progression. Cluster 2 showed a gradual increase from SP through AK followed by a sharp rise into cSCC and contained `LY96`, `LAMP3`, `CLEC9A`, and `CCL19`, suggesting emergence of a mature migratory DC state with TLR4-accessory signaling. Cluster 1 was biphasic, with low expression in SP, a transient dip in SD, and progressive amplification through AK into cSCC; it contained `CD274`, `IRF1`, `ISG15`, `CXCL9`, `IDO1`, `HAVCR2`, `CASP4`, and `IL1RN`. This late-emerging program combines IFN/JAK–STAT-driven `PD-L1` induction with IDO1-mediated tryptophan catabolism and `TIM-3` co-expression, indicating a multilayered immunosuppressive DC phenotype that consolidates in established cSCC. A complementary endothelial signal reinforced this picture: endothelial cluster 1 declined toward cSCC and contained `CD274`, `PDCD1LG2`, `IL6`, `STAT1`, `IRF1`, and `CIITA`, suggesting loss of vascular antigen-presenting capacity including MHC class II regulation, while endothelial cluster 2 increased toward cSCC with `ENTPD1`, `TBK1`, `IFI30`, and `CCL8`, consistent with a shift toward an activated vascular immune-interface state.

#### T cell consequences: loss of effector function and checkpoint expansion

T cell trajectories showed corresponding changes in effector and regulatory programs. Cluster 1 declined progressively into cSCC and contained `TRAF6`, `TNF`, and `CD40LG`, indicating erosion of costimulatory and effector signaling. Cluster 3 increased gradually throughout the disease continuum and contained `FOXP3`, `CTLA4`, `TIGIT`, `ENTPD1`, and `CXCL13`, consistent with progressive expansion of regulatory and checkpoint-high T cell states from early disease stages onward; the progressive accumulation of `CXCL13` across this cluster also marks exhausted tumor-reactive CD8^+^ T cells and is associated with tertiary lymphoid structure formation. Cluster 2 showed low to intermediate expression across SP, SD, and AK before a sharp late rise in cSCC, and contained `PDCD1`, `CCL5`, `GNLY`, and `GZMB`, consistent with a PD-1^+^ cytotoxic tumor-reactive program. Together, these T cell trajectories define a “hot but exhausted” microenvironment in established cSCC, in which cytotoxic activity is present but ultimately restrained by co-expanding regulatory programs.

#### Keratinocyte immune-interface reprogramming

Keratinocyte trajectories identified two opposing programs that together reflect staged epithelial reprogramming beginning at the SD stage. Cluster 1 initiated at SD and accelerated through AK into cSCC, with coordinated upregulation of MHC class II antigen-presentation genes (`HLA-DRA`, `HLA-DRB1`, `HLA-DPA1`, `HLA-DPB1`), immunoproteasome component `PSMB9`, IFN-stimulated genes (`IFITM3`, `IFI6`, `IFI27`), and TLR4-associated inflammatory mediators (`S100A8`, `S100A9`, `CASP4`), consistent with acquisition of a non-professional APC phenotype under inflammatory conditions. Cluster 2 showed a progressive decline to a nadir at AK with partial recovery in cSCC and contained homeostatic IFN*γ* signaling components (`IFNGR1`, `STAT3`, `IRF1`), physiological NF-*κ*B activity (`NFKB1`), and epithelial maintenance ligands (`AREG`, `HBEGF`). Loss of `AREG` and `HBEGF` may amplify DAMP release and TLR4 engagement, establishing a feed-forward loop between epithelial homeostatic failure and innate inflammatory activation. Both keratinocyte programs are already initiated at the SD stage, reinforcing SD as a biologically active window for topical immunoprevention from the epithelial compartment as well as the immune compartment.

Collectively, these trajectories support a model of staged immune remodeling in which TLR4- associated myeloid activation and stromal inflammatory signaling emerge as early as SD, IFN-driven PD-L1 induction in dendritic cells consolidates through AK into cSCC, and a “hot but exhausted” T cell microenvironment characterized by simultaneous cytotoxic and regulatory checkpoint programs becomes established in invasive disease.

## 3 Discussion

The development of effective topical immunoprevention strategies for cSCC represents an urgent and actionable clinical need. Preclinical models have established the blockade of `PD-L1`/`PD-1` and `TLR4` as promising intervention targets; however, the stage at which these pathways become biologically active in human skin has not yet been defined in human tissue. The present findings directly address this gap. By mapping the single-cell transcriptomic landscape across the SP–SD–AK–cSCC continuum, we demonstrate that meaningful activation of immune checkpoints and innate inflammatory remodeling are detectable well before invasive malignancy, and key signals emerge as early as sun-damaged skin. This has immediate implications for the design of topical immunoprevention trials that should be timed prior to the development of AK. The early appearance of dendritic cells in the SD stage, absent from the skin of SP, is among the most clinically actionable findings of this study. Dendritic cells are professional antigen-presenting cells and central orchestrators of adaptive immune priming, and their emergence in chronically sun-damaged tissue suggests that the immune system already responds to UV-induced cellular damage prior to histologically confirmed dysplasia. These dendritic cells co-expressed `PD-L1` (`CD274`), consistent with an immunoregulatory rather than purely activating phenotype. `PD-L1` expression by dendritic cells has been shown to attenuate T cell activation and blunt antitumor immunity, and IFN/JAK–STAT/`IRF1` signaling, a well-established inducer of `PD-L1`, was also upregulated in this population [Garcia-Diaz et al., 2017, Oh et al., 2020, Peng et al., 2020]. Together, these observations suggest that topical `PD-L1` blockade applied at the sun-damaged stage could restore dendritic cell-mediated immune priming before an immunosuppressive niche is fully established, consistent with the rationale underlying the topical small-molecule `PD-L1` blockade seen by other research [Dickinson et al., 2024].

The myeloid signal associated with `TLR4`, which is most clearly manifested in macrophages, provides a complementary and mechanistically distinct rationale for early intervention. Macrophage expression of `CD14`, `LY96` (`MD-2`) and `CD86` progressively increased throughout the trajectory from SD to cSCC, consistent with progressive activation of the `TLR4` receptor complex throughout carcinogenesis. `LY96/MD-2` is an essential co-receptor that enables `TLR4` to respond to both exogenous and endogenous danger signals, including `HMGB1`, which was simultaneously upregulated in fibroblasts across the same disease trajectory. This `HMGB1`–`TLR4` axis has been proposed as a mechanism that links UV-induced tissue damage to chronic innate inflammatory signaling [Kim et al., 2013], and pharmacological `TLR4` antagonism with resatorvid has previously been shown to block solar UV-induced skin tumorigenesis in murine models [Blohm-Mangone et al., 2018]. Chronic UV exposure causes keratinocyte apoptosis and necrosis, releasing `HMGB1` and other damage-associated molecular patterns (DAMPs) in the extracellular space, where they engage the `TLR4`–`MD-2` (`LY96`) receptor complex on resident myeloid cells [Johnson et al., 2013, Bald et al., 2014]. The progressive co-upregulation of `HMGB1` in fibroblasts and `LY96` in macrophages observed here is consistent with a self-sustaining DAMP-driven innate inflammatory cycle that accumulates with each successive stage of UV-induced tissue injury. Current data extend this biology into human tissue, providing single-cell evidence that `TLR4`-associated myeloid activation accumulates progressively throughout the human carcinogenesis continuum and is not restricted to the tumor microenvironment.

The findings of T cells add an important layer of complexity to the immunoprevention picture. The late cSCC-associated expansion of `PD-1`-positive cytotoxic T cells (`PDCD1`, `GNLY`, `GZMB`) in conjunction with regulatory and checkpoint-high T cell programs (`FOXP3`, `CTLA4`, `TIGIT`, `ENTPD1`, `CXCL13`) suggests that at the time of invasive malignancy, the immune microenvironment has undergone substantial counter-regulatory remodeling. The T-cell cluster 1 showed a progressive decline into cSCC and contained `TRAF6`, `TNF`, and `CD40LG`, indicating loss of effector and costimulatory T-cell programs during malignant progression. Critically, the trajectory of this effector T cell decline warrants closer stage-level examination: if the loss of `TRAF6` and `CD40LG` costimulatory signaling is already initiated at the SD or AK stage rather than at the point of cSCC, this would imply that the functional window for preserving rather than restoring effector T cell competence is earlier than the AK stage, and would further strengthen the rationale for immunoprevention intervention in clinically SD but histologically normal skin.

The co-expansion of effector and suppressive T cell states in cSCC mirrors patterns observed in other solid tumors and likely reflects a partially active but ultimately restrained antitumor immune response [Duhen et al., 2018, De Simone et al., 2021]. The relative scarcity of these populations in AK compared to cSCC reinforces the notion that the AK-to-cSCC transition represents a critical inflection point and that earlier intervention at the SD or AK stage may be necessary to prevent the establishment of this suppressive T cell niche. The marked enrichment of plasma and B cells in cSCC (in the UMAP), combined with progressive upregulation of `CXCL13`, a key B cell-recruiting chemokine, across T cell populations further suggests that humoral immune remodeling accompanies the lymphoid changes. Although formal trajectory analysis of plasma and B cell populations was beyond the scope of this study, due to limited representation across earlier disease stages, the role of B cell infiltration and potential TLS-associated humoral activity in the cSCC immune landscape warrants dedicated investigation in future larger cohorts. The progressive increase in `CXCL13` expression from SD through AK and in cSCC further marks exhausted tumor-reactive CD8^+^ T cells and is associated with B-cell recruitment and the formation of tertiary lymphoid structures (TLS) in solid tumors [Workel et al., 2019, Chen et al., 2025]. Whether `CXCL13`-driven TLS formation at the SD and AK stages represents a productive antitumor immune response worth preserving, or an early marker of chronic immune dysregulation that ultimately facilitates exhaustion remains an open question; resolving this distinction is important for immunoprevention as interventions that inadvertently disrupt nascent TLS could impair rather than restore local antitumor immunity.

The stromal and endothelial findings, while secondary to the myeloid and lymphoid signals, suggest that the immunoprevention microenvironment extends beyond the immune cells. Progressive upregulation of `CXCL9`, `CXCL10`, and `IL6` by fibroblasts along with `IFN`-responsive genes indicates that stromal cells contribute to the chemokine environment that recruits and shapes immune populations throughout carcinogenesis. The late upregulation of `IDO1` and `TAP1` in cSCC suggests that stromal immunoregulatory mechanisms may contribute to immune evasion in the invasive stage. The co-expression of `IDO1` and `CD274` (`PD-L1`) in dendritic cell cluster 1 suggests that the immunosuppressive phenotype of DCs in cSCC may involve multiple layers, combining `PD-L1`-mediated T cell inhibition with `IDO1`-driven tryptophan catabolism. This pattern is consistent with dual immunosuppressive mechanisms reported in other solid tumors and raises the possibility that combined `IDO1` inhibition and `PD-L1` blockade could be worth exploring in advanced cSCC lesions [Platten et al., 2019].

Endothelial remodeling toward an activated immune-interface state (`ENTPD1`, ;;TBK1, `CCL8`) in cSCC is consistent with vascular reprogramming in the tumor microenvironment and may represent an additional layer of immune regulation at the blood vessel interface. Endothelial cell trajectories revealed divergent changes in immune-interface programs. Cluster 1, which declined during progression to cSCC, expressed `CD274` (`PD-L1`), `PDCD1LG2` (`PD-L2`), `IL6`, `STAT1`, `IRF1`, and `CIITA`. The progressive loss of `CIITA`, a regulator of MHC class II antigen presentation, suggests that vascular antigen-presenting capacity is reduced in the cSCC microenvironment. This may represent an additional immune evasion mechanism operating in parallel with checkpoint molecule upregulation, potentially impairing T cell priming or reactivation at the vascular interface [Zheng et al., 2025]. In contrast, cluster 2 increased toward cSCC and was enriched for `ENTPD1`, `TBK1`, `IFI30`, and `CCL8`, consistent with a shift toward an activated vascular immune-interface state that may facilitate immune cell trafficking into the tumor. Although these stromal and vascular programs are unlikely to be the primary intervention targets, they can influence the efficacy and penetration of topically applied agents and merit consideration in the design of delivery strategies. The stromal inflammatory programs identified here are consistent with the broader concept of skin inflammaging described by Furman et al., in which chronic low-grade inflammation driven by senescent cell accumulation and SASP factor secretion reshapes the skin microenvironment in ways that promote both aging and carcinogenesis [Furman et al., 2025].

The keratinocyte trajectory analysis extends this picture beyond the immune compartment to the epithelial compartment itself. The opposing keratinocyte trajectories, progressive acquisition of an `IFN`-driven immune-interface program alongside simultaneous suppression of homeostatic `IFN`-*γ* responsiveness and epithelial maintenance signaling, suggests a staged epithelial reprogramming that mirrors the immune dysregulation observed across myeloid, dendritic cell, and T cell compartments. Both programs are already initiated at the SD stage, reinforcing SD and AK as biologically active stages for immunoprevention not only from the immune side but from the epithelial side as well. These data support a model in which the optimal window for topical immunoprevention directed by `PD-L1`/`PD-1` and `TLR4` spans the stages of SD to AK early enough so that the immunosuppressive niche is not yet fully established, but late enough so that the relevant immune populations and pathway activity are already present and detectable. Recent clinical evidence directly supports this window: immune checkpoint inhibitor therapy has been shown to significantly reduce AK burden in high-risk populations, with mean AK counts declining from 47 to 14 at 12 months [Cox et al., 2025], and topical calcipotriol plus 5-fluorouracil immunotherapy at the AK stage reduced SCC incidence through induction of tissue-resident memory T cell responses [Rosenberg et al., 2019]. This cellular roadmap, with key signals emerging at the SD stage, supports the rationale for topical rather than systemic checkpoint blockade as a targeted immunoprevention strategy.

Whether the PD-L1/TLR4 transcriptional signature identified here predicts progression risk from SD skin to AK or cSCC remains an important question for prospective cohort studies. A critical practical consideration for translating these findings into topical immunoprevention trials is whether small-molecule `PD-L1` and `TLR4` antagonists can achieve sufficient dermal penetration to engage the immune populations identified here, the majority of which, including macrophages, T cells, and the migratory dendritic cell populations identified in this study, reside predominantly within the dermis. Prior work by Dickinson et al. demonstrated that topical small-molecule `PD-L1` blockade inhibits UV-induced inflammatory signaling in murine skin [Dickinson et al., 2024], explicit pharmacokinetic characterization of dermal drug concentrations in human skin at each carcinogenesis stage will be essential to confirm that therapeutically relevant exposures can be achieved at the sites of immune remodeling identified in the present study.

This study has several limitations that should be considered in interpreting the findings. The sample size, while powered for primary cell population detection, is modest, and cSCC cases are not matched to SP-AK trajectory samples, excluding within-individual comparisons in the malignant stage. Furthermore, the unmatched cSCC group differed from the paired SP–SD–AK cohort with respect to age, sex distribution, and prior cSCC burden (Table 1), and the potential contribution of these demographic differences to the transcriptional signals observed in cSCC cannot be evaluated in this experiment. Additionally, while the co-expression patterns identified across cell types suggest coordinated intercellular and intracellular signaling, the present study did not apply formal ligand-receptor communication analysis such as `CellChat` or `NicheNet`; future studies incorporating such approaches would help determine whether these inferred interactions reflect functional intercellular signaling rather than transcriptional co-occurrence, and would more precisely define the cellular communication networks that govern immune remodeling across the carcinogenesis continuum. Replication in larger prospectively collected cohorts with paired spatial transcriptomic and protein-level validations will be important to confirm the stage-specific signals identified here and to determine whether the immune microenvironment features observed translate to predictors of progression risk at the patient level.

**Table 1:** Sample Demographics.

| Characteristic | Category | Paired SP-SD-AK<br>(n = 8) | cSCC<br>(n = 8) | Total<br>(n = 16) |
| --- | --- | --- | --- | --- |
| Sex | Male | 5 (62%) | 7 (88%) | 12 (75%) |
|  | Female | 3 (38%) | 1 (12%) | 4 (25%) |
| Age (years) | Median (Range) | 73 (62–81) | 82 (64–87) | 80 (62–87) |
| Fitzpatrick Skin Type | Type I | 5 (62%) | 3 (38%) | 8 (50%) |
|  | Type II | 3 (38%) | 2 (25%) | 5 (31%) |
|  | Unknown | 0 (0%) | 3 (38%) | 3 (19%) |
| No. of Previous BCC | Median (Range) | 1 (0–5) | 0 (0–5) | 0 (0–5) |
| No. of Previous cSCC | Median (Range) | 1 (0–3) | 3 (0–29) | 2 (0–29) |
| No. of Previous Melanoma | Median (Range) | 0 (0–2) | 0 (0–2) | 0 (0–2) |
| Previous Actinic Keratosis | Yes | 8 (100%) | 8 (100%) | 16 (100%) |

*Values are n (%) for categorical variables and median (range) for continuous variables.*
*Abbreviations: SP = sun-protected skin; SD = sun damaged skin; AK = actinic keratosis; cSCC = cutaneous squamous cell carcinoma; BCC = basal cell carcinoma.*

**Table 2:** Fuzzy C-Means Trajectory Summary.

| Cell type | Cluster | Trajectory | Key genes |
| --- | --- | --- | --- |
| Dendritic cells | 1 | Low SP/SD,<br>increase AK→cSCC | CD274, IRF1, ISG15,<br>CXCL9, IDO1, HAVCR2 |
|  | 2 | Increasing to cSCC | LY96, LAMP3, CLEC9A,<br>CCL19, HAVCR2 |
|  | 3 | Decreasing to cSCC | IL1B, TNF, CLEC10A,<br>CXCL8 |
| Macrophages<sup>†</sup> | 2 | Increasing<br>SP→cSCC | CD14, LY96, CD86, CCL18 |
| T cells | 1 | Decreasing to cSCC | TRAF6, TNF, CD40LG |
|  | 2 | Late increase in<br>cSCC | PDCD1, CCL5, GNLY, GZMB |
|  | 3 | Gradual increase<br>SP→cSCC | FOXP3, CTLA4, TIGIT,<br>ENTPD1, CXCL13 |
| Fibroblasts | 1 | Increasing<br>SP→cSCC | HMGB1, IL6, STAT1,<br>CXCL9, CXCL10 |
|  | 2 | Late increase in<br>cSCC | LY96, IDO1, MX1, TAP1,<br>CXCL13 |
|  | 3 | Decreasing to cSCC | TICAM2, IRAK4, TRAF6 |
| Endothelial cells | 1 | Decreasing to cSCC | CD274, PDCD1LG2, IL6,<br>STAT1, IRF1, CIITA |
|  | 2 | Increasing to cSCC | ENTPD1, TBK1, IFI30,<br>CCL8 |
| Keratinocytes | 1 | Increasing<br>SD→cSCC | HLA-DRA, HLA-DRB1,<br>HLA-DPA1, HLA-DPB1,<br>PSMB9, IFITM3, S100A8,<br>S100A9, CASP4 |
|  | 2 | Decreasing to nadir<br>at AK, partial<br>recovery in cSCC | IFNGR1, STAT3, IRF1,<br>NFKB1, AREG, HBEGF |

<sup>†</sup>Macrophage clusters 1 and 3 had insufficient cell representation across SP, SD, and AK to yield interpretable trajectories (see Figure 1). SP = sun-protected; SD = sun-damaged; AK = actinic keratosis; cSCC = cutaneous squamous cell carcinoma; APC = antigen-presenting cell; TLS = tertiary lymphoid structure; DAMP = damage-associated molecular pattern.

## 4 Materials and methods

### 4.1 Sampling frame and study design

The study was approved by an institutional review board disclosed in the Author Details document, and written informed consent was obtained from all participants. Skin biopsies were obtained from eight individuals with matched sun-protected skin (SP, n=8), sun-damaged skin (SD, n=8), and actinic keratoses (AK, n=8) samples, together with eight unmatched cutaneous squamous cell carcinoma (cSCC, n=8). The handling of tissues followed the Gurtner lab skin tissue protocol [Chen et al., 2021]. At the time of sample acquisition, skin samples were split for fresh single-cell RNA sequencing (scRNA-seq) analysis and a small section for formalin-fixed and paraffin-embedded (FFPE) and hematoxylin and eosin staining (H&E) to confirm diagnosis. All tissues were dissociated within 2 hours of receipt. If the sample had multiple dissociation vials, they were combined into one tube at this point. Cells were counted using a 1:1 ratio of cells to trypan blue on a Countess II Automated Cell Counter (Invitrogen, AMQAX1000), and delivered to an institutional genetic core for single-cell RNAseq on the 10X Genomics Chromium platform at a final concentration of 1000 cells/*µ*L. The tissue samples were clinically annotated with the corresponding detailed phenotypic information, i.e., the skin phototype, the degree of SD assessed by a standardized scale [McKenzie et al., 2011], the total number of AKs by anatomical location, the history of skin cancer and the risk level of cSCC as shown in Table 1.

### 4.2 Raw data preprocessing

Raw single-cell RNA sequencing files (FASTQ) were processed using the standard Cell Ranger pipeline to obtain gene-by-cell UMI count matrices [Zheng et al., 2017]. All of the following analyses were performed in `R` (v4.5.1; [R Core Team, 2025]) unless otherwise specified. The `Seurat` package (v5.4.0; [Hao et al., 2023]) was used for quality control and downstream analyses. Filtering was performed separately for each sample using the raw RNA counts. Cells were retained if they had fewer than 50,000 total UMIs, between 300 and 8,000 detected genes, and a proportion of mitochondrial transcripts below the overall sample median plus two standard deviations. Mitochondrial percentage was calculated using genes with prefixes `MT`or `mt-`. To mitigate ambient RNA contamination, the samples were further processed with `decontX` (v1.8.0; [Yin et al., 2025]) and the decontaminated assay was used for all primary analyses. In addition, putative doublets were identified using `scDblFinder` (v1.24.10; [Germain et al., 2022]), and only singlets were retained for all downstream analyses.

### 4.3 Cell type annotation

Cell type annotation was performed using `CellTypist` (v1.7.1; [Xu and Teichmann, 2022]) in `Python` (v3.9.20; [Van Rossum and Drake, 2009]) with the `Adult_Human_Skin` v1 reference model, followed by manual consolidation into broad biologically interpretable lineages. These lineages included keratinocytes, fibroblasts, endothelial cells, macrophages, dendritic cells, T cells, natural killer (NK) cells, plasma/B cells, mast cells, melanocytes, pericytes, glandular epithelium, and Schwann cells. Low-abundance populations with limited representation across conditions were retained for global visualization but excluded from selected pseudobulk differential expression, pathway, and trajectory analyses when sample coverage was inadequate to ensure stable inference.

### 4.4 Targeted trajectory clustering over cancer progression

For the pathway-level trend analysis shown in Figure 2, we evaluated selected genes from the `PD-L1`/`PD-1` checkpoint axis and the `TLR4`-associated innate inflammatory pathway. Pseudobulk counts were normalized using the trimmed mean of M values (TMM) method, followed by transformation to logarithmic counts per million (log-CPM) [Robinson et al., 2010, Robinson and Oshlack, 2010]. To assess monotonic expression changes across the ordered SP→SD→AK→cSCC sequence, the Jonckheere–Terpstra test was applied to pseudobulk log-CPM values for each gene. Benjamini–Hochberg correction was used to control the false discovery rate across genes tested in this pathway-focused trend analysis.

For the targeted trajectory clustering shown in Figure 3, we performed fuzzy c-means clustering using a focused panel of genes representing six functional categories: `PD-L1`/`PD-1` checkpoint regulation, `TLR4`-associated innate inflammatory signaling, IFN/NAK–STAT-mediated `PD-L1` induction, antigen presentation/APC maturation, chemokine-mediated immune recruitment, and immunoregulatory/tissue-remodeling support. Stage-wise expression was summarized using the median log-CPM across pseudobulk samples from each cell type and disease stage, yielding a four-stage trajectory over SP, SD, AK, and cSCC for each gene. Genes were retained if they were present after preprocessing and showed nonzero variance across stages. Gene expression values were standardized to have mean 0 and standard deviation 1 before clustering.

**Figure 3.**
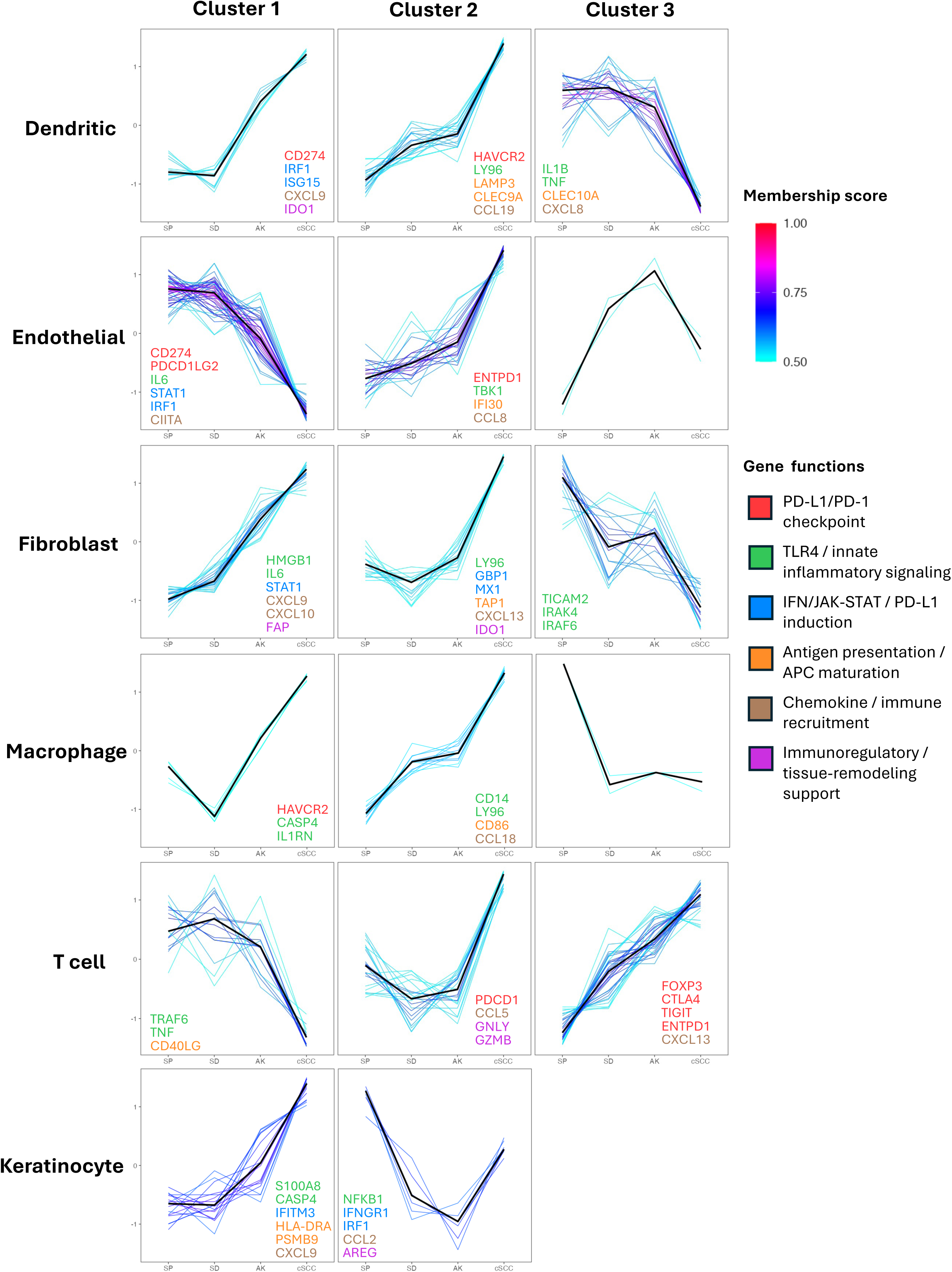
Pathway targeted fuzzy c-means clustering revealed the expression trajectory of targeted genes across SP, SD, AK, and cSCC. Pathway-focused fuzzy c-means clustering was performed across SP, SD, AK, and cSCC. Rows represent selected cell types, and columns represent clusters. Colored curves show standardized gene trajectories, and black curves show cluster centroids. Labeled genes indicate representative genes from the targeted functional categories. Unlabeled panels indicate that too few targeted genes passed the representative membership threshold.

To identify both monotonic and non-monotonic trajectory patterns, we applied soft clustering using the Mfuzz package (v2.70.0; [Kumar and Futschik, 2007]). Unlike hard clustering methods, which assign each gene to a single cluster, soft clustering allows a gene to belong to multiple clusters with varying degrees of membership, a framework particularly well suited for gene expression data where genes may participate in multiple biological processes and exhibit overlapping trajectory patterns. Let *ỹ_gj_* denote the standardized expression value for gene *g* (*g* = 1*, …, G*) in disease stage *j* (*j* = 1*, …, J*). Mfuzz implements fuzzy c-means clustering by minimizing the objective function

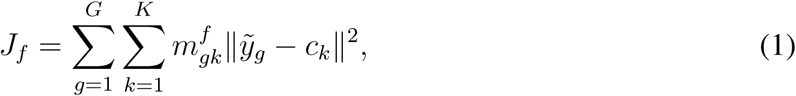

where *K* is the number of clusters, *ỹ_g_* = (*ỹ_g_*_1_, …, *ỹ_gJ_*)^⊤^ is the standardized expression profile for gene *g*, *c_k_*is the centroid of cluster *k*, *m_gk_* ∈ [0, 1] is the membership value of gene *g* in cluster *k*, and *f >* 1 is the fuzzification parameter controlling the degree of overlap among clusters [Futschik and Carlisle, 2005]. The optimal fuzzification parameter *f* was estimated separately for each cell type using the `mestimate()` function in Mfuzz, yielding the following values: dendritic cells, *f* = 3.45; endothelial cells, *f* = 3.32; fibroblasts, *f* = 3.27; macrophages, *f* = 3.82; T cells, *f* = 3.34; keratinocytes, *f* = 4.82. Because the analysis included only four ordered stages, we used three clusters per cell type to capture the major interpretable trajectory patterns — increasing, decreasing, and stage-specific or non-monotonic — while avoiding over-fragmentation. Genes were assigned to clusters according to their maximum membership score, and only genes with a membership score ≥ 0.5 were retained as representative for interpretation. Cell types with insufficient representation across SP, SD, and AK were excluded from trajectory analysis when sample coverage was inadequate to ensure stable inference; for macrophages, only one cluster yielded interpretable gene-level results due to sparse representation in pre-malignant stages (see Figure 1).

## Author Contributions

Conceptualization: RW, RP, BJL, XS, CCL, SED Data Curation: RW, RP, BJL, XS Formal Analysis: RW, RP Funding Acquisition: BJL, CCL, SED, GTW Investigation: RW, RP, DS, SC, CCL, GTW, SED, BJL, XS Methodology: RW, RP, BJL, XS Project Administration: BJL, CCL, SED, XS Resources: SC, CCL, GTW, SED, BJL, XS Software: RW, RP Supervision: BJL, XS Validation: RW, RP, BJL, XS Visualization: RW, RP, XS Writing – Original Draft: RW, RP, BJL, XS Writing – Review and Editing: RW, RP, DS, SC, CCL, GTW, SED, BJL, XS

## Data and Code Availability Statement

Single-cell RNA sequencing data generated in this study will be deposited in the Gene Expression Omnibus before publication. Processed objects and metadata will be made available through GEO-linked supplemental repository. The analysis code will be made available through a public GitHub repository before publication.

## Ethics Statement

All participants provided written, informed consent, and this protocol was approved by the institutional review board at the University of Arizona (Protocol Title: Skin Cancer Prevention Program Biorepository; Protocol Number: 1200000229).

## Artificial Intelligence Statement

During the preparation of this manuscript, the authors used ChatGPT (OpenAI, GPT-5.5) and Claude (Anthropic, Opus 4.8) to assist with grammar and language editing and to generate example R and Python code for data analysis. The prompts included requests for grammar revision of manuscript text and generation of R and Python code implementing statistical analyses described by the authors. All AI-generated text and code were carefully reviewed, edited, and verified by the authors. In particular, all code was independently evaluated, tested, and validated before use. The authors take full responsibility for the content of this manuscript and the accuracy of all analyses and results.

## Acknowledgments

The research reported in this publication was supported by the National Cancer Institute of the National Institutes of Health under Award Numbers P30 CA023074, P01 CA229112, and UG3 CA290443. The content is solely the responsibility of the authors and does not necessarily represent the official views of the National Institutes of Health. Additional support was provided by the University of Arizona Cancer Center Skin Institute through philanthropic donations, and by the University of Arizona Office of Research and Partnerships Accelerate for Success program.

## Abbreviations

AK: actinic keratosis
cSCC: cutaneous squamous cell carcinoma
DC: dendritic cell
FFPE: formalin-fixed paraffin-embedded
GSEA: gene set enrichment analysis
IFN: interferon
NK: natural killer
NMSC: nonmelanoma skin cancer
PD-1: programmed cell death protein 1
PD-L1: programmed death-ligand 1
scRNA-seq: single-cell RNA sequencing
SD: sun-damaged skin
SP: sun-protected skin
TLR4: Toll-like receptor 4
UMAP: uniform manifold approximation and projection.

## References

Fabrizio Antonangeli, Ambra Natalini, Marina Chiara Garassino, Antonio Sica, Angela Santoni, and Francesca Di Rosa. Regulation of PD-L1 expression by NF-*κ*B in cancer. Frontiers in Immunology, 11:584626, 2020. doi: 10.3389/fimmu.2020.584626. PMID: 33324403.

Peter Bailey, Rachel A. Ridgway, Patrizia Cammareri, Mairi Treanor-Taylor, et al. Driver gene combinations dictate cutaneous squamous cell carcinoma disease continuum progression. Nature Communications, 14:5211, 2023. doi: 10.1038/s41467-023-40822-9.

Tobias Bald, Thomas Quast, Jennifer Landsberg, Meri Rogava, Nicole Glodde, Dorys Lopez-Ramos, Judith Kohlmeyer, Stefanie Riesenberg, Debby van den Boorn-Konijnenberg, Cornelia Hömig-Hölzel, Raphael Reuten, Benjamin Schadow, Heike Weighardt, Daniela Wenzel, Iris Helfrich, Dirk Schadendorf, Wilhelm Bloch, Marco E. Bianchi, Claire Lugassy, Raymond L. Barnhill, Manuel Koch, Bernd K. Fleischmann, Irmgard Förster, Wolfgang Kastenmüller, and Thomas Tüting. Ultraviolet-radiation-induced inflammation promotes angiotropism and metastasis in melanoma. Nature, 507:109–113, feb 2014. doi: 10.1038/nature13111.

Kimberly Blohm-Mangone, Nancy B. Burkett, Shah Mohammad Tahsin, Paul B. Myrdal, Aronganaa Aodengtuya, Billy Ho, Sreevidya Rayala Uppalapati, Janice Berlin, Clara Curiel-Lewandrowski, Hari K. Pal, Craig A. Elmets, Mohammad Athar, and Sally E. Dickinson. Pharmacological TLR4 antagonism using topical resatorvid blocks solar UV-induced skin tumorigenesis in SKH-1 mice. Cancer Prevention Research, 11(5):265–278, 2018. doi: 10.1158/1940-6207.CAPR-17-0349.

J. T. Chen, S. J. Kempton, and V. K. Rao. The economics of skin cancer: An analysis of medicare payment data. Plastic and Reconstructive Surgery Global Open, 4:e868, 2016. doi: 10.1097/GOX.0000000000000826.

K. Chen et al. Disrupting biological sensors of force promotes tissue regeneration in large organisms. Nature Communications, 12:5256, 2021. doi: 10.1038/s41467-021-25410-z.

Yulu Chen, Y. Wu, Z. Zhao, L. Wen, M. Wu, D. Song, Q. Zeng, Y. Liu, G. Yan, and G. Zhang. Retrospective study on the correlation between CXCL13, immune infiltration, and tertiary lymphoid structures in cutaneous squamous cell carcinoma. PeerJ, 13:e19398, 2025. doi: 10.7717/peerj.19398. URL 10.7717/peerj.19398. PMID: 40352278.

Magdalena Ciąyńska, Irmina Olejniczak-Staruch, Dorota Sobolewska-Sztychny, Joanna Narbutt, Małgorzata Skibićska, and Aleksandra Lesiak. Ultraviolet radiation and chronic inflammation— molecules and mechanisms involved in skin carcinogenesis: A narrative review. Life, 11(4):326, 2021. doi: 10.3390/life11040326. PMID: 33917793.

Charlotte Cox, Susan Brown, Euan Walpole, and, et al. Immune checkpoint inhibitors in field cancerization and keratinocyte cancer prevention. JAMA Dermatology, 161(4):383–390, apr 2025. doi: 10.1001/jamadermatol.2024.5750. URL 10.1001/jamadermatol.2024.5750.

Vincent D. Criscione, Martin A. Weinstock, Mark F. Naylor, Claudia Luque, Melody J. Eide, and Stephen F. Bingham. Actinic keratoses: Natural history and risk of malignant transformation in the Veterans Affairs Topical Tretinoin Chemoprevention Trial. Cancer, 115(11):2523–2530, June 2009. doi: 10.1002/cncr.24284.

Vincenzo De Falco, Stefania Napolitano, Renato Franco, Federica Zito Marino, Luigi Formisano, Daniela Esposito, Gabriella Suarato, Rossella Napolitano, Alfonso Esposito, Francesco Caraglia, et al. Overexpression of CCL-20 and CXCL-8 genes enhances tumor escape and resistance to cemiplimab, a programmed cell death protein-1 (PD-1) inhibitor, in patients with locally advanced and metastatic cutaneous squamous cell carcinoma. OncoImmunology, 13(1): 2388315, 2024. doi: 10.1080/2162402X.2024.2388315.

Marco De Simone, Alessandra Arrigoni, Giulia Rossetti, Paolo Gruarin, Valentina Ranzani, Claudia Politano, Raoul J. P. Bonnal, Enrico Provasi, Maria L. Sarnicola, Isabella Panzeri, et al. Transcriptional landscape of human tissue lymphocytes unveils uniqueness of tumor-infiltrating t regulatory cells. Communications Biology, 4:1012, 2021. doi: 10.1038/s42003-021-02537-9.

Sally E. Dickinson and Georg T. Wondrak. TLR4 in skin cancer: From molecular mechanisms to clinical interventions. Molecular Carcinogenesis, 58(7):1086–1093, 2019. doi: 10.1002/mc.23016. PMID: 30892799.

Sally E. Dickinson, Prajakta Vaishampayan, Jana Jandova, Yuchen (Ella) Ai, Viktoria Kirschnerova, Tianshun Zhang, Valerie Calvert, Emanuel Petricoin, H-H. Sherry Chow, Chengcheng Hu, Denise Roe, Ann Bode, Clara Curiel-Lewandrowski, and Georg T. Wondrak. Inhibition of uv-induced stress signaling and inflammatory responses in skh-1 mouse skin by topical small-molecule pd-l1 blockade. JID Innovations, 4(2): 100255, March 2024. ISSN 26670267. doi: 10.1016/j.xjidi.2023.100255. URL https://linkinghub.elsevier.com/retrieve/pii/S2667026723000826.

Thomas Duhen, Rebekka Duhen, Ryan Montler, Jake Moses, Tarsem Moudgil, Noel F. C. C. de Miranda, Cheri P. Goodall, Tiffany C. Blair, Bernard A. Fox, Jason E. McDermott, et al. Co-expression of cd39 and cd103 identifies tumor-reactive cd8 t cells in human solid tumors. Nature Communications, 9:2724, 2018. doi: 10.1038/s41467-018-05072-0.

David Furman, Johan Auwerx, Anne-Laure Bulteau, et al. Skin health and biological aging. Nat. Aging, 5(7):1195–1206, July 2025. doi: 10.1038/s43587-025-00901-6. URL 10.1038/s43587-025-00901-6.

Matthias E Futschik and Bronwyn Carlisle. Noise-robust soft clustering of gene expression timecourse data. Journal of Bioinformatics and Computational Biology, 3(04):965–988, 2005.

Angel Garcia-Diaz, David S. Shin, Beatriz Homet Moreno, Jorge Saco, Helena Escuin-Ordinas, Gabriela A. Rodriguez, Jesse M. Zaretsky, Lili Sun, Willy Hugo, Xueqiao Wang, Giuseppe Parisi, Claudio P. Saus, Daniel Y. Torrejon, Thomas G. Graeber, Begonya Comin-Anduix, Siwen Hu-Lieskovan, Robert Damoiseaux, Roger S. Lo, and Antoni Ribas. Interferon receptor signaling pathways regulating pd-l1 and pd-l2 expression. Cell Reports, 19(6):1189–1201, 2017. doi: 10.1016/j.celrep.2017.04.031.

Pierre-Luc Germain, Aaron Lun, Carlos Garcia Meixide, Will Macnair, and Mark D. Robinson. Doublet identification in single-cell sequencing data using scdblfinder. f1000research, 2022. doi: 10.12688/f1000research.73600.2.

Gery P Guy Jr, Steven R Machlin, Donatus U Ekwueme, and K Robin Yabroff. Prevalence and costs of skin cancer treatment in the US, 2002- 2006 and 2007- 2011. American Journal of Preventive Medicine, 48(2):183–187, 2015.

Yuhan Hao, Tim Stuart, Madeline H Kowalski, Saket Choudhary, Paul Hoffman, Austin Hartman, Avi Srivastava, Gesmira Molla, Shaista Madad, Carlos Fernandez-Granda, and Rahul Satija. Dictionary learning for integrative, multimodal and scalable single-cell analysis. Nature Biotechnology, 2023. doi: 10.1038/s41587-023-01767-y. URL 10.1038/s41587-023-01767-y.

Yuan Jiang, Yueyuan Zheng, Yuan-Wei Zhang, Shuai Kong, Jinxiu Dong, Fei Wang, et al. Reciprocal inhibition between TP63 and STAT1 regulates antitumor immune response through interferon-*γ* signaling in squamous cancer. Nature Communications, 15(1):2479, 2024. doi: 10.1038/s41467-024-46785-9. URL 10.1038/s41467-024-46785-9.

Kelly E. Johnson, Brian C. Wulff, Tatiana M. Oberyszyn, and Traci A. Wilgus. Ultraviolet light exposure stimulates HMGB1 release by keratinocytes. Archives of Dermatological Research, 305(9):805–815, nov 2013. doi: 10.1007/s00403-013-1401-2.

Seung Kim, So-Young Kim, John P. Pribis, Michael Lotze, Kevin P. Mollen, Robert Shapiro, Patricia Loughran, Melanie J. Scott, and Timothy R. Billiar. Signaling of high mobility group box 1 through toll-like receptor 4 in macrophages requires cd14. Journal of Immunology, 190 (12):6454–6462, 2013. doi: 10.4049/jimmunol.1203478.

Lokesh Kumar and Matthias E. Futschik. Mfuzz: a software package for soft clustering of microarray data. Bioinformation, 2(1):5–7, 2007. doi: 10.6026/97320630002005.

Bonnie LaFleur, Clara Curiel-Lewandrowski, Edgar Tapia, Joel Parker, Lisa White, H-H Sherry Chow, and Andrew P. South. Characterizing dermal transcriptional change in the progression from sun-protected skin to actinic keratosis. Journal of Investigative Dermatology, 143(7):1299–1302.e3, July 2023. doi: 10.1016/j.jid.2022.12.021. URL 10.1016/j.jid.2022.12.021.

Wiltrud Lederle, Sofia Depner, Sabine Schnur, Eva Obermüller, Nicola Catone, Alexandra Just, Norbert E. Fusenig, and Margareta M. Mueller. IL-6 promotes malignant growth of skin SCCs by regulating a network of autocrine and paracrine cytokines. International Journal of Cancer, 128(12):2803–2814, 2011. doi: 10.1002/ijc.25621. PMID: 20726000.

Zhenlin Li, Fangqi Lu, Fujin Zhou, Dekun Song, Lunhui Chang, Weiying Liu, Guorong Yan, and Guolong Zhang. From actinic keratosis to cutaneous squamous cell carcinoma: the key pathogenesis and treatments. Frontiers in Immunology, 16: 1518633, January 2025. ISSN 1664-3224. doi: 10.3389/fimmu.2025.1518633. URL https://www.frontiersin.org/articles/10.3389/fimmu.2025.1518633/full.

N. E. McKenzie et al. Development of a photographic scale for consistency and guidance in dermatologic assessment of forearm sun damage. Archives of Dermatology, 147(1):31–36, 2011. doi: 10.1001/archdermatol.2010.392.

Michael R. Migden et al. Pd-1 blockade with cemiplimab in advanced cutaneous squamous-cell carcinoma. New England Journal of Medicine, 2018. doi: 10.1056/NEJMoa1805131.

A. M. Mousa et al. Immune checkpoints and cellular landscape of the tumor microenvironment in non-melanoma skin cancer. Cells, 13(19):1615, 2024. doi: 10.3390/cells13191615.

Soyoung A. Oh, Dai-Chen Wu, Jeanne Cheung, Armando Navarro, Huizhong Xiong, Rafael Cubas, Klara Totpal, Henry Chiu, Yan Wu, Laetitia Comps-Agrar, et al. Pd-l1 expression by dendritic cells is a key regulator of t-cell immunity in cancer. Nature Cancer, 1:681–691, 2020. doi: 10.1038/s43018-020-0075-x.

Qi Peng, Xuewen Qiu, Zhonglin Zhang, Siyuan Zhang, Yanhua Zhang, Yefeng Liang, Jiao Guo, Hong Peng, Ming Chen, Y. X. Fu, et al. Pd-l1 on dendritic cells attenuates t cell activation and regulates response to immune checkpoint blockade. Nature Communications, 11:4835, 2020. doi: 10.1038/s41467-020-18570-x.

Michael Platten, Ellen A. A. Nollen, Ute F. Röhrig, Francesca Fallarino, and Christiane A. Opitz. Tryptophan metabolism as a common therapeutic target in cancer, neurodegeneration and beyond. Nature Reviews Drug Discovery, 18(5):379–401, may 2019. doi: 10.1038/s41573-019-0016-5. Review.

R Core Team. R: A Language and Environment for Statistical Computing. R Foundation for Statistical Computing, Vienna, Austria, 2025. URL https://www.R-project.org/.

Mark D. Robinson and Alicia Oshlack. A scaling normalization method for differential expression analysis of rna-seq data. Genome Biology, 11:R25, 2010. doi: 10.1186/gb-2010-11-3-r25.

Mark D. Robinson, Davis J. McCarthy, and Gordon K. Smyth. edger: a bioconductor package for differential expression analysis of digital gene expression data. Bioinformatics, 26(1):139–140, 2010. doi: 10.1093/bioinformatics/btp616.

Howard W Rogers, Martin A Weinstock, Steven R Feldman, and Brett M Coldiron. Incidence estimate of nonmelanoma skin cancer (keratinocyte carcinomas) in the us population, 2012. JAMA Dermatology, 151(10):1081–1086, 2015.

Abby R. Rosenberg, Mary Tabacchi, Kenneth H. Ngo, Michael Wallendorf, Ilana S. Rosman, Lynn A. Cornelius, and Shadmehr Demehri. Skin cancer precursor immunotherapy for squamous cell carcinoma prevention. JCI Insight, 4(6):e125476, mar 2019. doi: 10.1172/jci.insight.125476. URL 10.1172/jci.insight.125476.

Rosalba Salcedo, Christophe Cataisson, Uzma Hasan, Nabiha Yusuf, and Giorgio Trinchieri. MyD88 and its divergent toll in carcinogenesis. Trends in Immunology, 34(8):379–389, 2013. doi: 10.1016/j.it.2013.03.008.

Samuel Schepps, Jonathan Xu, Henry Yang, Jenna Mandel, Jaanvi Mehta, Julianna Tolotta, Nicole Baker, Volkan Tekmen, Neda Nikbakht, Paolo Fortina, Ignacia Fuentes, Bonnie LaFleur, Raymond J. Cho, and Andrew P. South. Skin in the game: a review of singlecell and spatial transcriptomics in dermatological research. Clinical Chemistry and Laboratory Medicine, 62(10):1880–1891, September 2024. doi: 10.1515/cclm-2023-1245. URL 10.1515/cclm-2023-1245.

Mohammad Asif Sherwani, Carlos Alberto Mier Aguilar, Charlotte McRae, Gelare Ghajar-Rahimi, Aisha Anwaar, Ahmed Omar Jasser, Ariq Chandra, Hui Xu, and Nabiha Yusuf. MyD88 plays an important role in UVB-induced suppression of dendritic cell activity, T cell function, and cutaneous immune response. International Journal of Molecular Sciences, 26(19):9361, 2025. doi: 10.3390/ijms26199361. URL 10.3390/ijms26199361.

Selin Tokez, Loes Hollestein, Marieke Louwman, Tamar Nijsten, and Marlies Wakkee. Incidence of multiple vs first cutaneous squamous cell carcinoma on a nationwide scale and estimation of future incidences of cutaneous squamous cell carcinoma. JAMA Dermatology, 156(12):1300– 1306, 2020.

Zewen K. Tuong, Andrew Lewandowski, Jennifer A. Bridge, Jazmina L. G. Cruz, Miko Yamada, Duncan Lambie, Richard Lewandowski, Raymond J. Steptoe, Graham R. Leggatt, Fiona Simpson, et al. Cytokine/chemokine profiles in squamous cell carcinoma correlate with precancerous and cancerous disease stage. Scientific Reports, 9(1):17754, 2019. doi: 10.1038/s41598-019-54435-0.

Prajakta Vaishampayan, Clara Curiel-Lewandrowski, and Sally E. Dickinson. Review: PD-L1 as an emerging target in the treatment and prevention of keratinocytic skin cancer. Molecular Carcinogenesis, 62(1):52–61, 2023. doi: 10.1002/mc.23464.

Guido Van Rossum and Fred L. Drake. Python 3 Reference Manual. CreateSpace, Scotts Valley, CA, 2009. ISBN 1441412697.

Hagma H. Workel, Joyce M. Lubbers, Roland Arnold, Thalina M. Prins, Pieter van der Vlies, Kim de Lange, David van der Meer, G. Bea A. Wisman, Marcel A. T. M. van Vugt, Hans W. Nijman, and Marco de Bruyn. A transcriptionally distinct CXCL13+CD103+CD8+ t-cell population is associated with b-cell recruitment and neoantigen load in human cancer. Cancer Immunology Research, 7(5):784–796, May 2019. doi: 10.1158/2326-6066.CIR-18-0517. URL 10.1158/2326-6066.CIR-18-0517.

Chuan Xu and Sarah A. Teichmann. Cross-tissue immune cell analysis reveals tissue-specific features in humans. Science, 376(6594), 2022. doi: 10.1126/science.abl5197. URL https://www.science.org/doi/10.1126/science.abl5197.

Tao Xu, Zhongzhi Wang, Hao Wu, Yuanjie Zhu, Liangzhe Wang, Siyu Wu, Ti Fu, Xiaolie Wang, Guotai Yao, Shengli Li, Yi Jin, and Jianghan Chen. Single-cell transcriptomic landscape of keratinocyte transformation from actinic keratosis to skin carcinoma. iScience, 28(12):114169, dec 2025. doi: 10.1016/j.isci.2025.114169. Epub 2025 Nov 25. eCollection 2025 Dec 19.

Guorong Yan et al. Unraveling the landscape of non-melanoma skin cancer through single-cell rna sequencing technology. Frontiers in Oncology, 14:1500300, 2024. doi: 10.3389/fonc.2024.1500300.

Hanlin Yin, Ning Pu, Qiangda Chen, Jicheng Zhang, Guochao Zhao, et al. Gut-derived lipopolysaccharide remodels tumoral microenvironment and synergizes with PD-L1 checkpoint blockade via TLR4/MyD88/AKT/NF-*κ*B pathway in pancreatic cancer. Cell Death & Disease, 12(11):1033, 2021. doi: 10.1038/s41419-021-04293-4. PMID: 34718325.

Yuan Yin, Masanao Yajima, and Joshua Campbell. decontX: Decontamination of single cell genomics data, 2025. URL https://bioconductor.org/packages/decontX. R package version 1.8.0.

Grace X. Y. Zheng, Jessica M. Terry, Phillip Belgrader, Paul Ryvkin, Zachary W. Bent, Ryan Wilson, Solongo B. Ziraldo, Tobias D. Wheeler, Geoff P. McDermott, Junjie Zhu, Mark T. Gregory, Joe Shuga, Luz Montesclaros, Jason G. Underwood, Donald A. Masquelier, Stefanie Y. Nishimura, Michael Schnall-Levin, Paul W. Wyatt, Christopher M. Hindson, Rajiv Bharadwaj, Alexander Wong, Kevin D. Ness, Lan W. Beppu, H. Joachim Deeg, Christopher McFarland, Keith R. Loeb, William J. Valente, Nolan G. Ericson, Emily A. Stevens, Jerald P. Radich, Tarjei S. Mikkelsen, Benjamin J. Hindson, and Jason H. Bielas. Massively parallel digital transcriptional profiling of single cells. Nature Communications, 8(1):14049, January 2017. ISSN 2041-1723. doi: 10.1038/ncomms14049. URL https://www.nature.com/articles/ncomms14049.

Tianyu Zheng, Xinran Yu, Caihong Yu, Wangting Xu, Zhuoyang Fan, Yongjie Zhou, Changyu Li, Juncheng Wan, Chaoqiao Jin, Xuran Jin, Wen Zhang, Zhiping Yan, Peng Luo, Bufu Tang, and Xudong Qu. Tumor-associated endothelial cells in tumor immune escape and immunotherapy: multifaceted roles and treatment approaches. Biomarker Research, 14(1):10, dec 2025. doi: 10.1186/s40364-025-00883-y. Review.

W. Zou et al. Single-cell sequencing identifies the malignant progression and intercellular communication in actinic keratosis, squamous cell carcinoma in situ, and cutaneous squamous cell carcinoma. eLife, 2023. doi: 10.7554/eLife.85270. URL https://elifesciences.org/articles/85270.

